# Sequence engineering at non-motif modulator residues yields a peptide that effectively targets a single PDZ protein in a disease-relevant cellular context

**DOI:** 10.1101/2025.09.25.678608

**Authors:** Jeanine F. Amacher, Patrick R. Cushing, Lars Vouilleme, Sierra N. Cullati, Bin Deng, Scott A. Gerber, Prisca Boisguerin, Dean R. Madden

## Abstract

**Highlights:**

– PDZ domains are critical components of many cellular, disease-relevant systems
– Overlapping specificity complicates selective targeting of PDZ interaction networks
– We engineered a single-PDZ peptide inhibitor using non-motif preferences
– Mass-spectrometry based experiments confirm single-PDZ binding
– Our inhibitor increases chloride efflux in a cystic fibrosis cellular model

PDZ interaction networks are finely-tuned products of evolution. These widespread binding domains recognize short linear motifs (SLiMs), usually at the C-terminus of their interacting partners, and are involved in trafficking and signaling pathways, the formation of tight junctions, and scaffolding of the post-synaptic density of neurons, amongst other roles. Typically, a single PDZ domain binds multiple targets; conversely, each PDZ-binding protein engages several PDZ domains, dependent on cellular conditions. Historical PDZ binding motifs rely on two key positions for binding. However, previous insights on modulator, or non-motif, selectivity preferences reveal that these limited motifs are insufficient to describe PDZ-mediated interactomes, consistent with the observation that the degree of promiscuity is much more limited than predicted by defined binding classes. Here, we use these principles to engineer and test a peptide-based inhibitor capable of interacting with a single PDZ domain-containing protein in a disease-relevant cellular system. We first interrogate a previously developed sequence selective for cystic fibrosis transmembrane conductance regulator (CFTR)-Associated Ligand (CAL), one of five PDZ domains known to bind the CFTR C-terminus, probing for off-target PDZ partners. Once identified, we use parallel biochemical and structural refinement to eliminate these interactions and introduce a CAL PDZ inhibitor with unprecedented PDZ domain selectivity. We test and verify specificity using relevant cellular PDZ target networks in a mass spectrometry-based approach. Our resultant selective inhibitor enhances chloride efflux when applied to polarized patient bronchial epithelial cells, as well as confirms that engineering an effectively single-PDZ peptide is possible when modulator preferences are applied.

## Introduction

Complex logistics pathways and scaffolds in the cell are often coordinated by short linear motif (SLiM) targeting sequences [1–5]. An example of a SLiM-binding domain that plays a critical role in cellular regulation, including in the assembly and composition of complex macromolecular assemblies including neuronal synapses and epithelial junctions, is the PDZ domain [6,7]. Named for the first three discovered members of the family (<u>P</u>SD-95, <u>D</u>lg, and <u>Z</u>O-1), PDZ domains typically recognize peptides at the C-terminus of their binding partners [8–11]. These are ancient domains, present in bacteria, fungi, and animals, underscoring their critical importance in cellular networks [12–15].

PDZ proteins contain one or more these domains, with up to 13 in a single human protein and over 20 in a protein from our closest non-metazoan ancestor, the choanoflagellate *Monosiga brevicollis* [11,15]. Overall, 272 PDZ domains have been found in 184 proteins in the human proteome, all with a shared structural fold and similar binding mode, constituting the largest collection of protein-protein interaction domains [11,16]. PDZ domains were initially grouped into three classes (I-III), based on alignment of SLiMs for a given domain that often revealed motif preferences at only two residues: the P^0^ (the most C-terminal residue, preceded by P^-1^ etc.) and P^-2^ positions; however, early on these binding motifs were known to be insufficient to describe cellular PDZ interaction networks. [17–19]. Indeed, this set of classifications predicts more than 10,000 PDZ binding sequences in the human proteome, and often more than 1,000 for a single domain, suggesting either highly degenerate recognition or the presence of cryptic affinity determinants [20]. Certain PDZ domains can also interact with internal motifs, although the stereochemistry of ligand recognition remains similar [21,22].

More detailed biochemical analyses later led to the proposal of much more differentiated binding motifs for many PDZ domains, each dependent on up to seven residues in the target sequence [23]. However, in contrast to the original classification, the 16 newer binding classes (1a-h, 2a-f, 3a, 4a) predict only 12 potential PDZ-compatible C-termini proteome-wide, excluding almost all known PDZ:partner interactions. Thus, the predictive power of highly differentiated binding motifs is overly restrictive. Several recent computational and experimental approaches have thus been utilized to understand the PDZome in cellular systems [24–26]. Furthermore, recent work by ourselves and others reveals that some naturally-occurring PDZ target sequences are highly selective for a small number of PDZ domains, while others are more promiscuous. In particular, pathogenic proteins from several viruses, e.g., human papillomavirus (HPV), human immunodeficiency virus (HIV), influenza, and SARS-CoV-2, have evolved to hijack multiple cellular systems via binding to relatively large numbers of PDZ proteins [11,27–34]. For example, the holdup assay, which probed binding of a peptide mimicking the C-terminus of the HPV16 E6 oncoprotein to 209 PDZ domains, found that this sequence bound 20% of the human PDZome [33]. In contrast, endogenous PDZ target sequences tend to be more selective, binding a relatively small number of PDZ proteins [28]. These differences are attributed to *modulator* residues, or non-motif affinity determinants which co-determine peptide-binding affinity [15,20,28]. These observations suggest that target promiscuity can be tuned, but perhaps not eliminated, based on amino-acid characteristics at individual modulator positions.

Here, we test the hypothesis that optimization of a combination of modulator preferences would permit us to target a single PDZ domain with high specificity in a cell-based disease model. We chose to use the cystic fibrosis transmembrane conductance regulator (CFTR) and cystic fibrosis (CF) as a model system for these studies. The CFTR C-terminus is known to functionally bind several PDZ domains, including those of the Na^+^/H^+^ exchanger regulatory factor (NHERF) proteins and CFTR-associated ligand (CAL), which regulate its endocytic trafficking [35–37]. It may also interact with other proteins at their PDZ domains, including mast2 microtubule associated serine/threonine kinase (MAST205) and sorting nexin 27 (SNX27), although direct interactions have not be validated *in vitro* [38,39]. Our lab and others have previously shown that CAL PDZ (CALP) binding to CFTR negatively regulates its abundance at the epithelial apical membrane. CAL also affects the apical-membrane stability of the F508del variant (p.Phe508del) encoded by *F508del-CFTR* (mutation c.1521_1523delCTT in *CFTR*), the most prevalent mutation associated with CF disease [35–37,40–42]. Selective inhibition of the CALP-CFTR interaction increases WT and F508del CFTR levels at the plasma membrane and thus presents a promising therapeutic strategy for CF [40,43–48].

Utilizing an iterative peptide-array approach, we previously engineered a peptide inhibitor of the CALP-CFTR interaction, inhibitor of CAL (iCAL)36, a decameric peptide (sequence: ANSRWPTSII) that binds CALP, but not any of the NHERF1/2 PDZ domains [20,44,45,49]. However, it remained unclear how many additional cellular PDZ domains would also be affected. We now develop a pipeline based on a pull down in combination with mass spectrometry to probe PDZ-targets of a peptide sequence in a relevant cell-based system, capable of identifying interactions in the mid-micromolar range. We then extend this work to challenge the specificity of iCAL36 in a CF cell model. When a major off-target PDZ protein is identified, Tax-interacting protein 1 (TIP-1), we are able to use our peptide array-based engineering pipeline and structural studies to understand the basis of this interaction, and remove TIP-1 binding. We argue that because iCAL36 was designed to be specific for several CALP modulator preferences, this was not only doable, but able to be achieved in relatively few additional steps. Furthermore, these approaches can be utilized synergistically with other therapeutic innovations, e.g., developing PDZ “molecular sponge” nanoparticles aimed at a specific pathogen (here, SARS-CoV-2) or in creating PDZ domain/self-binding peptide fusion reagents [50,51]. Taken together, our data show that we can probe cell lysates of a relevant tissue type to identify and eliminate off-target interactors, and engineer a PDZ inhibitor that can uniquely target CAL despite extensive overlapping interaction networks.

## Results

### A mass-spectrometry based approach to detect PDZ partners of target sequences

With the exception of the *holdup assay* [31,33,52], most studies of PDZ binding have used individual PDZ domains as bait to identify high affinity sequences from phage libraries or peptide arrays, essentially screening a fraction of the given peptidome [20,23,53]. While powerful, the holdup assay also interrogates a large majority of the PDZome in an *in vitro* system, and does not reflect protein expression levels in a given cell or tissue type. Therefore, here, we flip the paradigm. In order to explore the spectrum of partners for a given peptide sequence across a cellular proteome, we immobilized biotinylated peptides and used them to capture interacting PDZ proteins (**Figure 1**). This approach has previously been used to identify PDZ targets of HPV E6 oncoprotein from various strains [54].

**Figure 1.**
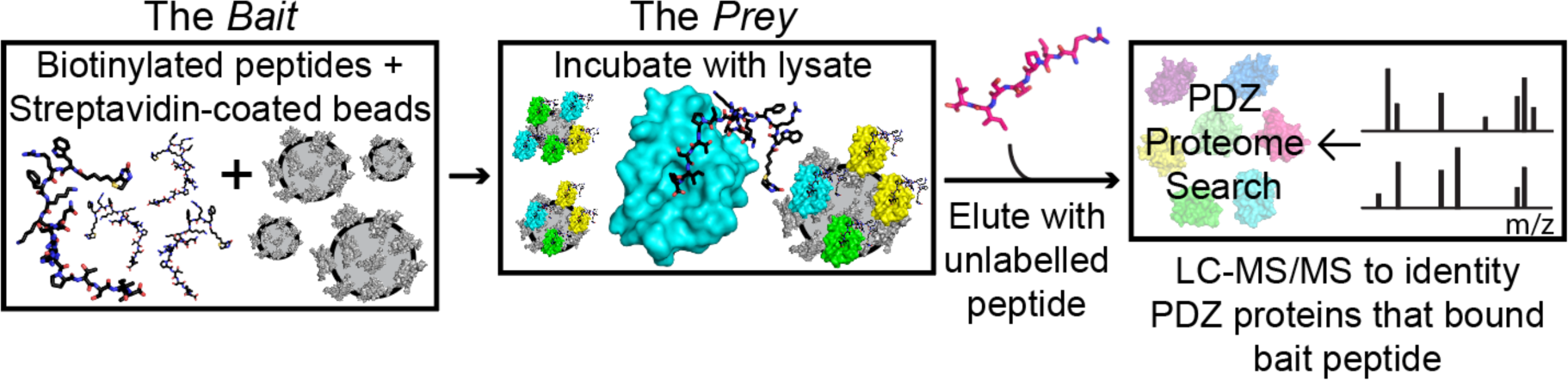
Detecting epithelial PDZ:peptide interactions by mass spectrometry. Experimental set-up of the pull-down experiment. The structure of CALP:BT-L-iCAL36 complex was determined (PDB ID 4Q6S), and is shown as a reference (also see **Table 2 and Figure S3**).

We performed pull-down experiments using biotinylated peptides bound to streptavidin-coated magnetic beads to capture interactors in filtered whole-cell lysate from polarized CFBE41o^−^ cystic fibrosis bronchial epithelial cells expressing *F508del* (CFBE-ΔF cells), as described in detail in Material and Methods. We then analyzed interacting proteins by tandem mass spectrometry (LC-MS/MS), in order to obtain an unbiased survey of the peptides’ individual epithelial interactomes. There are several challenges inherent to this approach. Physiologically relevant PDZ:peptide interactions often exhibit relatively weak affinities, with *K*_D_ values that can exceed 100 μM [1,11]. Since PDZ:peptide affinities are thought to have relatively high off-rates, a result of their reversible nature [55], a non-equilibrium assay is at particular risk of losing low-affinity partners. Furthermore, the relative protein expression levels of PDZ proteins in a given cell or tissue type is largely unknown.

To address these challenges and to maximize sensitivity in our pull-down protocol, we included several considerations in our protocol. First, we established the minimum number of rapid wash steps necessary to avoid non-specific signals. In addition, in order to limit the detection of lysate peptides non-specifically interacting with our bait, we used competitive elution with non-biotinylated cognate peptides to displace bound protein partners. We also tested a scrambled peptide (*BT*-SCR) as bait in parallel experiments to control for any remaining non-specific interactions; out of 184 human PDZ proteins, only five were identified, presumably not reflecting canonical PDZ interactions (see Methods in Supplementary material). Next, while we used SDS-PAGE analysis following the pull-down, we also captured the entire eluate and used unstained gels to resolve proteins for mass spectrometry analysis, which permitted identification of signals not seen with traditional silver-stained band identification (**Figure S1A**). Finally, we considered any PDZ protein identified with a relatively low number of unique peptides, ≥ 2, as a potential interactor, following up with additional biochemical experiments, e.g., Western blot, as needed.

For the initial characterization of our system, we tested two peptides that both exhibit Class I motifs (-X-S/T-X-Φ_COOH_, where X=any, Φ=hydrophobic). These included our model protein, CFTR (C-terminus:-VQDTRL_COOH_). The other was the viral protein HPV16 E6 (C-terminus:-RRETQL_COOH_), as previously tested in a similar experiment [54]. Both proteins bind multiple PDZ domains, but differ in their levels of promiscuity according to published scientific literature and interaction databases. While the HPV16 E6 peptide has been reported to bind >15 PDZ proteins [27,28,54,56–63], CFTR binds 5-7, as discussed earlier [28,38,39,64,65].

In our experiment, MS analysis identified at least two peptides for each of 13 validated PDZ domain-containing proteins for HPV16 E6 (**Figure 2A and Table S1**). We hypothesize that the failure to capture other previously-identified PDZ interactors, e.g., SNX27 and ZO-1/2, may be due to a variety of factors, including epithelial expression levels or protein chemistry. For example, while SNX27 has been identified in Caco-2 epithelial colon and Calu-3 human airway epithelial cells, in a bioRxiv preprint, we could find no reports looking at this protein in CFBE-ΔF cells [38,66]. Furthermore, ZO-1 expression is known to be disrupted in CFBE-ΔF cells, as compared to similar non-CF cell lines [67]. For *BT*-CFTR, we found only four PDZ proteins, including NHERF1/2/3 proteins, already known to interact with CFTR [65]. We also detected a small number of unique peptides for the PDZ protein, β-2-syntrophin (SNTB2, **Figure 2B and Table S1**), which is known to target the Ser/Thr kinase MAST205, a PDZ domain-containing protein that competes with CFTR-associated Ligand (CAL) for CFTR binding [39]. This may represent the detection of an indirect interaction, which is an additional feature of this approach.

**Figure 2.**
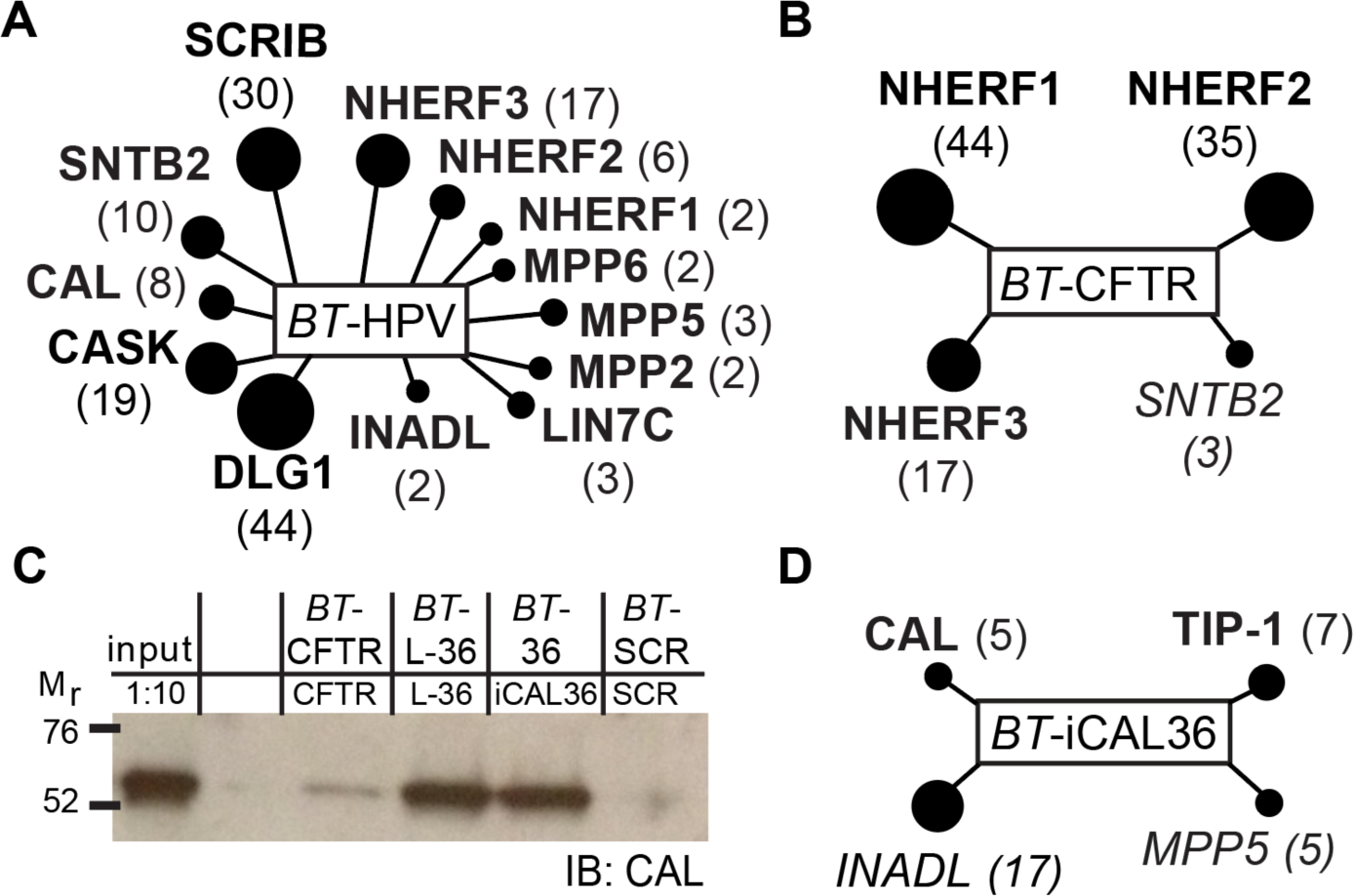
Differences in PDZ target promiscuity revealed. **(A-B,D)** Sequences identified by mass spectrometry of pull-down eluates using either BT-HPV (**A**) or BT-CFTR (**B**) peptides as bait. Sphere area is scaled to the number of peptides (in parentheses) identified via mass spectrometry (**Table S1**). Bold indicates a previously reported interaction. Italics indicate an interaction that has not been verified. **(C)** Immunoblot (IB) of pull-down eluates (50 µL; top: biotinylated bait sequences, bottom: elution peptides) reveal that CAL is robustly captured by iCAL36-based bait sequences and only weakly captured by BT-CFTR. The abbreviations and corresponding sequences of peptides used for capture and elution are described in Materials and Methods.

Overall, our data suggest that we successfully capture a majority of the previously reported interactors for the E6 and CFTR proteins. Furthermore, the discrepancy in the number of partners detected (e.g., 13 for E6 *versus* 4 for CFTR) most likely reflects the genuine difference in the promiscuity of the two sequences, rather than incomplete sampling of the population. Indeed, our recent work revealed that the combinatorial energetic contributions of several modulator residues in these sequences dramatically affects their respective number of PDZ targets [28]. These differences may reflect individual protein function, and are likely a product of evolution that could be shared in other viral versus endogenous sequences [28].

Our E6 and CFTR results were also used to calibrate the affinity threshold of our assay. We identified the CFTR associated ligand (CAL) in our *BT*-HPV16 E6 assay. CAL is known to bind *BT*-HPV with an affinity of 330 µM [20]. However, the CFTR C-terminus did not capture CAL, most likely reflecting the higher reported *K*_D_ values of 420-590 µM reported for this interaction [35,49] (**Figure 2B**). Based on these observations, our likely baseline detection threshold for our mass spectrometry-based approach is between 300 and 600 µM, a dramatic improvement over the reported 1-40 µM range typically accessible by phage display [53].

To compare the sensitivity of our MS-based assay with a traditional immunoblot for a known target, we repeated the pulldown procedure and assayed for the presence of CAL using an antibody. Although mass spectrometry did not detect CAL following capture with *BT*-CFTR, immunoblotting does, albeit as a weak band (**Figure 2C**). Thus, a highly specific antibody may provide slightly greater sensitivity than mass spectrometry, for a single, known target. In contrast, mass spectrometry can survey a much broader range of partners in parallel. Overall, our protocol is well-suited to assess peptide:PDZ selectivity across the epithelial proteome.

### Off-target PDZ binding of a CAL inhibitor engineered for limited selectivity

Our CFTR and HPV16 E6 results suggest that our pull-down with mass spectrometry-based assay is well-suited to explore PDZ specificity within a specific cell type. Thus, we next sought to interrogate our previously engineered CFTR inhibitor, iCAL36 [44,45]. Despite overlapping binding preferences among the PDZ domains that target CFTR, iCAL36 (ANSRWPTSII) successfully inhibits the PDZ domain of CAL, but not those of the NHERF proteins [45]. This is critical for the therapeutic targeting of CFTR as CAL facilitates the post-endocytic lysosomal degradation of CFTR, while the NHERF proteins stabilize CFTR at the apical membrane and enhance its chloride channel activity [36,40,41,68–71]. Because iCAL36 matches the frequently observed class I binding motif, however, it might interact with additional PDZ domains not tested in our original studies.

Indeed, using biotinylated *BT*-iCAL36, as our peptide bait, we captured not only CAL, but also three additional off-target PDZ proteins that we had not previously considered for negative design: TIP-1, InaD-like protein (InaDL) and MAGUK p55 subfamily member 5 (MPP5) (**Figure 2D**). TIP-1, InaDL, and MPP5 contain 1, 10, and 1 PDZ domains, respectively. TIP-1 has been previously implicated in protein trafficking pathways [72–74] and has an established preference for a P^-5^ Trp, as in the C-terminal sequence of β-catenin, <u>W</u>FDTDL [74]. Thus, we decided to investigate this interaction first, and verified it as an iCAL36 target in our pull-down assay by Western blot (**Figure S1B**).

We developed recombinant expression and purification protocols for the TIP-1 PDZ domain and monitored its interaction with a fluoresceinated iCAL36 peptide (*F\**-iCAL36) by fluorescence polarization (FP). Titration with *F\**-iCAL36 reveals a binding isotherm with a fitted *K*_D_ of 0.54 ± 0.04 µM. TIP-1 actually binds *F\**-iCAL36 2.5-fold more tightly than does CAL (*K*_D_ = 1.3 µM), and its submicromolar interaction places it at the high-affinity end of the spectrum of PDZ:peptide interactions (**Table 1**) [11,19,45]. TIP-1 binds *BT*-iCAL36 with an affinity of 2.4 ± 0.3 µM, which is similar to CAL (3.1 ± 1 µM) (**Table 1**). Taken together, our data suggest that *F\**-iCAL36 inhibits TIP-1 at least as potently as CAL. As a first step towards improved specificity, we therefore sought to design CAL inhibitors with selectivity expanded to exclude TIP-1.

**Table 1.** Fluorescence polarization binding affinities for PDZ domains with engineered peptides. The NHERF PDZ domains (N1P1, N1P2, N2P1, and N2P2) bind iCAL36 and L-iCAL36 peptides with *K*_i_ > 1000 μM, these were previously reported for iCAL36 [20,45]. All values for CALP, with the exception of *BT*-iCAL36 were previously reported [20,49].

| Peptide | Sequence | $K_i$ ($\mu\text{M}$) | |
| --- | --- | --- | --- |
|  |  | CALP | TIP-1 |
| iCAL36 | ANSR**W**PTSII | $22.6 \pm 8.0$ | $1.6 \pm 0.8$ |
| *BT*-iCAL36 | *BT*-ANSR**W**PTSII | $3.1 \pm 1.0$ | $2.4 \pm 0.3$ |
| F-iCAL36 | ANSR**F**PTSII | $110 \pm 20$ | $200 \pm 80$ |
| Y-iCAL36 | ANSR**Y**PTSII | $85 \pm 2$ | $390 \pm 15$ |
| L-iCAL36 | ANSR**L**PTSII | $68 \pm 20$ | $>1000$ |

### A stereochemical Achilles’ heel

As a basis for eliminating the off-target interaction with TIP-1, we investigated the roles of individual iCAL36 side chains using parallel structural and biochemical approaches. As mentioned above, the published structure of TIP-1 in complex with the β-catenin target peptide had revealed a deep pocket that accommodates the Trp residue at the P^-5^ position [74]. To confirm a similar binding mode for the P^-5^ Trp in iCAL36, and to visualize the stereochemistry of the interaction, we determined the crystallographic structure of TIP-1 bound to iCAL36 (**Figures S2, S3A**) at a resolution of 1.24 Å (**Table 2**). Like the previously reported structure of TIP-1 in complex with iCAL36 carrying a C-terminal Leu substitution, iCAL36_L_, the iCAL36 peptide adopts a canonical Class I PDZ binding mode (**Figure S3A)** [11,49,75], and as expected, the P^-5^ side chain is buried in a deep, hydrophobic pocket (**Figure 3A**).

**Figure 3.**
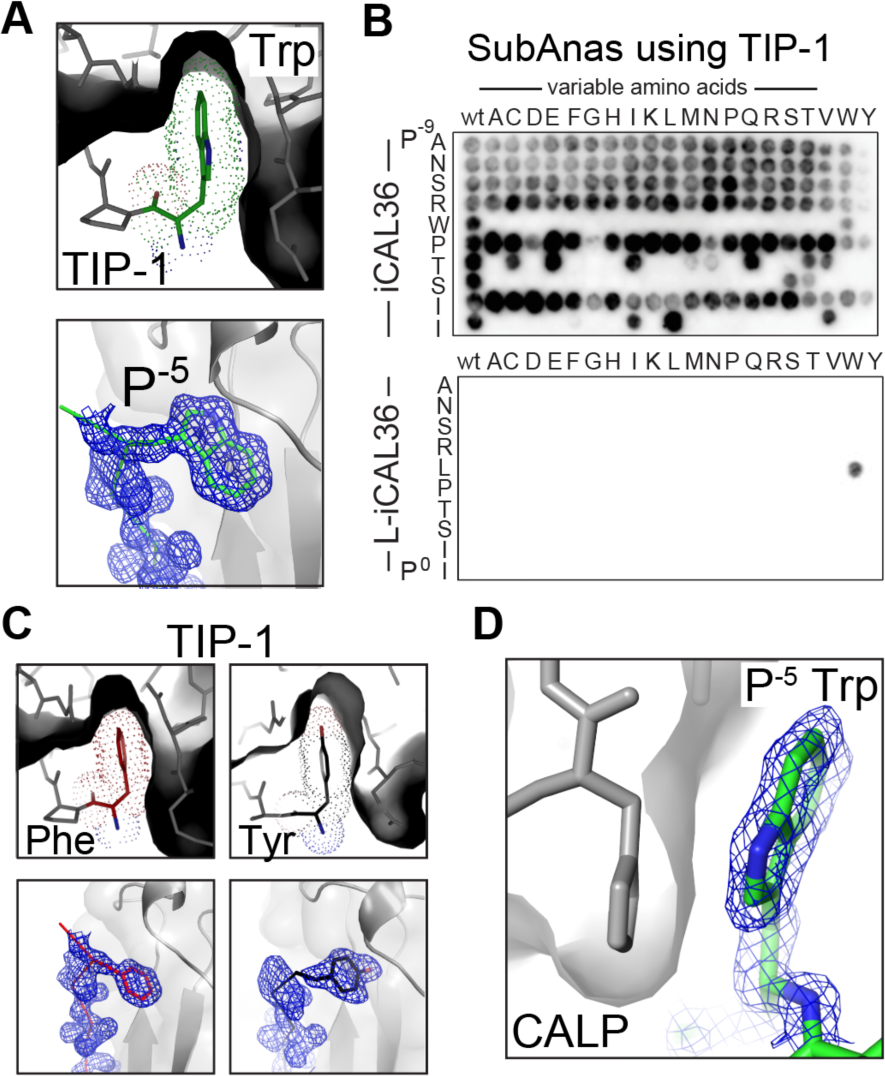
Targeting the TIP-1:iCAL36 Achilles’ heel. **(A,C)** Top: The van der Waals surface of the P^-5^ Trp-binding pocket is shown in cross section (grey surface), together with stick and dot-surface models of the bound iCAL36 Trp (green) (**A**), or (**C**) F-iCAL36 Phe (red) and Y-iCAL36 (black) side chains, illustrating the tight packing of the planar ring system. Bottom: Electron density, rendered at 1σ is shown for each of the 3 peptides (stick figure), respectively. TIP-1 is in gray surface and cartoon representation. **(B)** Binding of TIP-1 to SubAna arrays based on iCAL36 (top) or L-iCAL36 (bottom) reveals that TIP-1 binding requires a P^-5^ Trp. **(D)** Electron density, rendered at 1σ, is shown for the P^-5^ Trp in iCAL36 binding to CAL PDZ (PDB 4E34). The PDZ domain is in cross section (grey surface, sticks), together with a stick model of the bound iCAL36 (green sticks). Here, the superficial engagement of the P^-5^ Trp side chain with the surface of CALP is in contrast to the Trp-specific pocket in TIP-1 (highlighted in (**A**)).

**Table 2.**
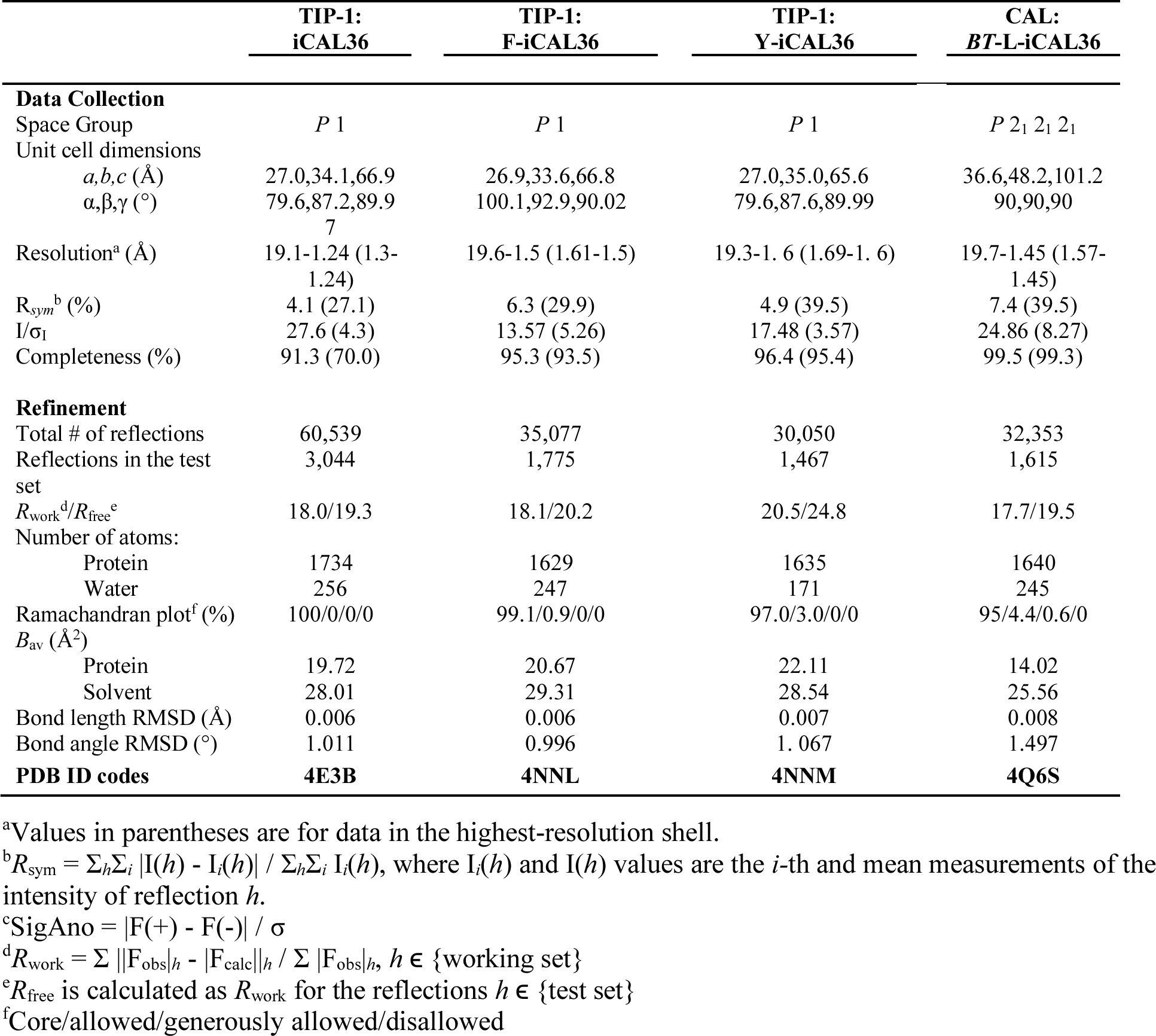
Data Collection and Refinement Statistics.

To identify a sequence that would retain selectivity for CALP, but no longer be able to bind TIP-1, we followed a combinatorial peptide array/FP counterscreening paradigm, similar to the one used in the initial design of iCAL36 [44,45]. A substitutional analysis (SubAna) array using iCAL36 as our starting sequence for TIP-1 fails to detect binding with sequences containing any residue other than Trp, suggesting that TIP-1 has a strong preference at P^-5^ (**Figure 3B, top**).

To test the extent of this preference given the hydrophobic binding pocket, we synthesized the closely related sequences F-iCAL36 (ANSR<u>F</u>PTSII) and Y-iCAL36 (ANSR<u>Y</u>PTSII). Crystal structures of each peptide in complex with TIP-1 at resolutions *d*_min_ ≤ 1.6 Å (**Table 2**) confirm that the substituted hydrophobic side chains are each bound in the same pocket that accommodates Trp (**Figure 3C**). Nevertheless, despite the excellent stereochemical complementarity to each planar aromatic conjugated ring system, binding affinity is weakened >300-fold in the presence of a P^-5^ Phe or Tyr (200 ± 80 µM for F-iCAL36 and 390 ± 15 µM for Y-iCAL36) (**Table 1**). This underscores the critical contribution of the P^-5^ Trp side chain to the affinity of iCAL36 for TIP-1.

To optimize CALP/TIP-1 discrimination without loss of selectivity against the NHERF domains, we recapitulated our analysis of decamer combinatorial libraries (CombLib), systematically varying the P^-4^ and P^-5^ residues within the framework of the iCAL36 sequence [45]. There were a number of combinations that bound CALP, but not TIP-1, with either a P^-4^ Pro or Gln (**Figure S4**). Of these, however, a P^-4^ Pro is required to avoid off-target interactions with the NHERF domains [45]. With this constraint, the highest CALP affinity is seen with a Cys at P^-5^, followed by Leu or Phe. Excluding Cys to avoid potential disulfide bond formation, we compared the binding preferences of F-iCAL36 and L-iCAL36 (ANSR**L**PTSII). As previously mentioned, whereas F-iCAL36 retains some affinity for TIP-1 *in vitro*, in FP displacement assays with TIP-1 and each of the NHERF domains, L-iCAL36 is indistinguishable from a DMSO control at peptide concentrations up to 1 mM. L-iCAL36 also exhibits slightly higher affinity for the CALP domain than F-iCAL36 does (68 ± 20 μM versus 110 ± 20 μM, respectively) [20].

Additional characterization confirmed the favorable affinity profile of L-iCAL36. First of all, a SubAna starting from the L-iCAL36 sequence shows that no additional substitution can restore affinity for TIP-1 PDZ, except for the Leu->Trp revertant (**Figure 3B, bottom panel**). This is in contrast to the CALP binding site, which in previous studies was shown to retain affinity for multiple P^-5^ substitutions, as previously shown, perhaps due to its more non-selective binding surface (**Figure 3D**) [20,45].

Finally, to assess the overall trade-off between affinity and selectivity among this target cluster, we determined the binding affinity of L-iCAL36 for purified NHERF1 and NHERF2 PDZ domains. Indeed, L-iCAL36 revealed no detectable binding to the NHERF1/2 PDZ domains, defined as *K*_D_ > 1 mM based on our assay conditions. Taken together, L-iCAL36 binds CALP with a free energy substantially more favorable than any of the other partners (ΔΔG^0^ = −2.5 kcal/mol). Thus, although L-iCAL36 binds CAL about 3-fold weaker than iCAL36, this has been parlayed into a 60-fold difference relative to the *K*_i_ of the PDZ domain with the next highest affinity (**Figure S5**). We can therefore define a thermodynamic ‘target specificity window’ for selective peptide inhibitors of CALP. This is in contrast to iCAL36, which favors binding of TIP-1 over CALP (ΔΔG^0^ = +1 kcal/mol) [44,45], and thus has no window of specificity for CALP.

Conversely, the SubAnas (**Figure 3B**) and CombLib (**Figure S4**) arrays underscore the differential sensitivity of CAL and TIP-1 to changes at P^-5^: while CALP retains affinity for multiple alternatives, TIP1 only demonstrates affinity for Trp. This behavior likely reflects the CALP-centric elaboration of the original iCAL36 sequence [45], in which affinity and specificity were optimized at each position. As a result, CALP can tolerate numerous individual substitutions, each with a relatively modest effect on overall binding affinity. In contrast, the adventitious interaction with TIP-1 is not nearly as robust to sequence modification, and is eliminated by most of the changes that are tolerated by CALP.

### A PDZ-selective inhibitor

Having succeeded in engineering selectivity against a primary off-target PDZ, we next wanted to test the ability of L-iCAL36 to bind other PDZ domains. We determined the crystal structure of CAL:*BT*-L-iCAL36 to confirm that the biotin tag does not interact with the PDZ domain for this peptide (**Table 2 and Figure S3B**), which it does not. Using our mass spectrometry-based approach, *BT-*L-iCAL36 robustly isolates only CAL from epithelial cells (**Figure 4A**). To increase our chances of detecting off-target interactors, we ran this experiment twice. CAL was detected in both iterations. In one experiment, we detected two peptides from NHERF2, and in another, two peptides from InaDL. Notably, L-iCAL36 no longer binds MPP5, our other iCAL36 off-target interaction (**Figure 2D**).

**Figure 4.**
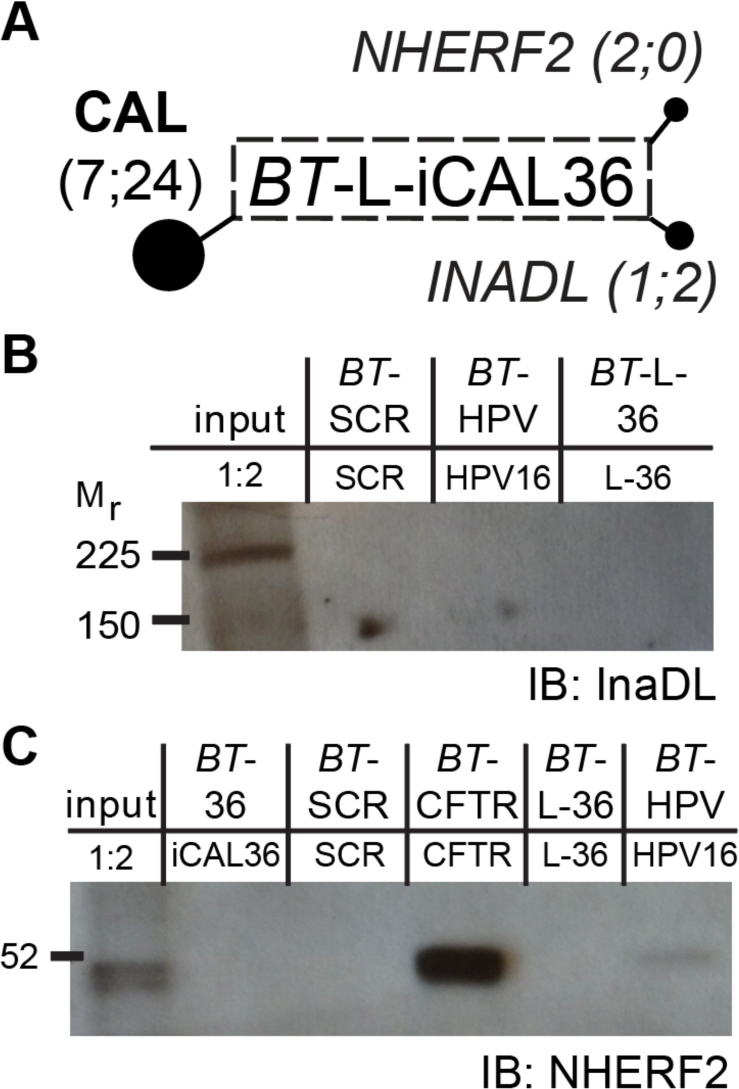
Endogenous PDZ networks are tuned by modulator preferences. **(A)** Results of pull-down, followed by mass spectrometry, experiments of the BT-L-iCAL36 peptide (dashed line indicates that the experiment was performed twice). Sphere area is scaled to the number of peptides (in parentheses) identified via mass spectrometry. Bold indicates a previously reported interaction. Italics indicate an interaction that had not been verified. **(B-C)** Immunoblots of pull-down eluates (50 µL; top: biotinylated bait sequences, bottom: elution peptides). Neither InaDL (**B**) nor NHERF2 (**C**) are detected in the BT-L-iCAL36 eluates. The abbreviations and corresponding sequences of peptides used for capture and elution are described in Materials and Methods.

To assess the extent of these interactions, we performed immunoblots of the eluates and were unable to identify either protein (**Figures 4B-C**). Whatever its exact affinity, InaDL also clearly interacts much more weakly with *BT*-L-iCAL36 (≤ 2 peptides) than with *BT*-iCAL36 (17 peptides; **Figure 2D**). Finally, FP analysis of L-iCAL36, similar to iCAL36, with either NHERF2 PDZ domain reveals no binding signal up to millimolar peptide concentrations. Both InaDL and NHERF2 are multi-PDZ proteins, and it is therefore possible that they leverage avidity effects in binding to the multivalent bead surface. Even with that advantage, their interactions remain at or beyond the threshold of detection. Overall, we conclude that L-iCAL36 strongly prefers binding to the single PDZ domain of CAL among the complete set of PDZ targets in this disease-relevant cellular system.

### F*-L-iCAL36 enhances CFTR-mediated Cl^-^ secretion

Finally, we exploited the high selectivity of L-iCAL36 to test the hypothesis that off-target interactions of iCAL36 might have contributed to the stabilization of F508del [44]. For these studies, we took advantage of the enhanced CALP selectivity of decapeptides carrying an N-terminal fluorescein moiety, which had been seen previously in comparison of the fluoresceinated (*K*_D_) and unlabeled (*K*_I_) affinities of iCAL36. We therefore synthesized an N-terminally fluoresceinated version of L-iCAL36 (*F\**-L-iCAL36) and tested it for binding against both CALP and TIP-1. In the context of the L-iCAL36 sequence, the addition of the N-terminal fluorescein moiety produces a 6-fold enhancement in CAL affinity, at 10.8 ± 0.2 µM. Conversely, the fluoresceinated peptide stills shows no appreciable binding to TIP-1 at the highest protein concentration tested, 1 mM.

Having validated the affinity profile of our fluoresceinated probe, we sought to determine whether *F*\*-L-iCAL36 is able to rescue F508del chloride-channel activity as efficiently as *F\**-iCAL36. In Ussing chamber measurements *F\**-iCAL36 and *F\**-L-iCAL36 were tested in head-to-head measurements for efficacy versus the scrambled control peptide, *F\**-SCR (**Figure 5**). Consistent with previous measurements, *F\**-iCAL36 increased the CFTR_inh_-172-sensitive short-circuit current (ΔI_sc_) by 10.7% (p = 0.0016; n=10).

**Figure 5.**
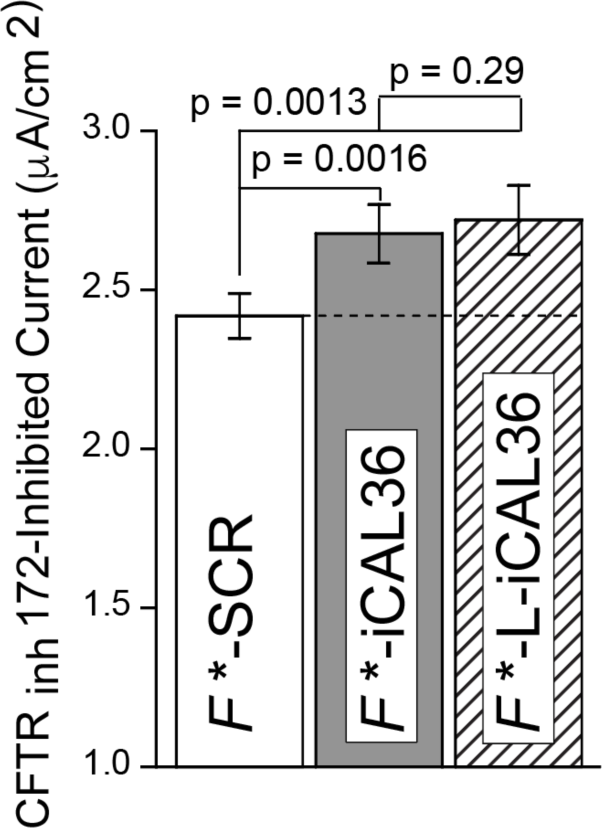
A globally selective CAL inhibitor increases CFTR-mediated Cl^-^ currents. Ussing chamber measurements confirm that L-iCAL36 retains efficacy for rescue of F508del CFTR. Polarized CFBE-ΔF cells were treated with F*-SCR, F*-iCAL36, or F*L-iCAL36. CFTR-specific short-circuit currents (ΔI_sc_) were measured following addition of CFTR_inh_-172. Mean ΔI_sc_ values are shown ± SEM (n=10).

Treatment of CFBE-ΔF cells with *F\**-L-iCAL36 yielded a 12.5% increase (p = 0.0013; n=10) in ΔI_sc_. Thus, *F\**-L-iCAL36 was at least as efficacious as *F*\*-iCAL36, suggesting that off-target interactions were not a substantial component of iCAL36-mediated chloride-channel rescue.

## Discussion

The design of inhibitors that selectively target PDZ domains is critical to our ability to dissect their individual regulatory contributions and may ultimately be a requirement for potential therapeutic applications. To overcome the well-known promiscuity of PDZ binding [9,11,18,35,76], we previously developed a peptide-engineering approach combining high-throughput peptide-array and high-precision FP affinity techniques to target only CALP among a cluster of PDZ domains that share affinity for the CFTR C-terminus [45].

Considering that our engineering efforts were targeted towards a specific cell type, we wanted to limit ourselves to only the relevant PDZ domains. Therefore, we chose to investigate specificity of our peptide inhibitors using a relevant cellular system, as opposed to available *in vitro* techniques, such as the holdup assay or a yeast two-hybrid approach [24,77–79]. While *in vitro* assays are more likely to identify irrelevant off-target interactors, due to their broader coverage of the PDZ interactome, here we demonstrate that a systematic survey of target specificity can be performed *post hoc*, utilizing pull-down assays to probe cell lysates of the relevant tissue type, and mass spectrometry to identify off-target interactors.

Our results reveal that the combinatorial diversity encoded in modulator preferences may thus help to explain the limited promiscuity that is observed in PDZ interaction networks, despite the highly degenerate motifs traditionally associated with PDZ binding. Indeed, although the CFTR C-terminus is seemingly compatible with the motifs of all Class I PDZ domains, our pull-down assay with *BT*-CFTR revealed only a few targets as compared to the HPV16 E6 C-terminus, which is known to be highly promiscuous. This likely reflects its pathophysiological importance as a broad-spectrum trafficking disruptor, and as mentioned previously, is due to combinatorial energetic contributions at modulator residues along the peptide-binding cleft [28].

This idea is further supported by comparisons of CALP and TIP-1 in our peptide array data. While it is difficult to relate SubAna spot intensities directly to affinity [80], at P^-5^, eight of the 19 replacements for Trp exhibited intensities at least 33% of the value for CAL binding to the iCAL36 starting sequence [20,45]. In contrast, every replacement for the Trp eliminates detectable binding for TIP-1 (**Figure 3B**).

When CAL *versus* TIP-1 preferences are evaluated in this way for each possible individual substitution at residues P^0^ through P^-5^ using SubAna array data, reported here and previously [45], 34 substitutions are identified that still bind CAL but not TIP-1 at 33% of wild-type iCAL36 intensity (blue boxes, **Figure 6**), compared with only three substitutions that retain TIP-1, but not CAL binding (yellow boxes, **Figure 6**). Considerable selectivity is seen even at relatively solvent-exposed side chains, such as P^-3^ and P^-4^ (**Figure 6**), underscoring the potential contributions of non-motif modulatory interactions along the full length of the peptide-binding cleft [20]. As a result, only a small sacrifice of CAL affinity was able to generate a substantial window of selectivity (**Figure S5**).

**Figure 6.**
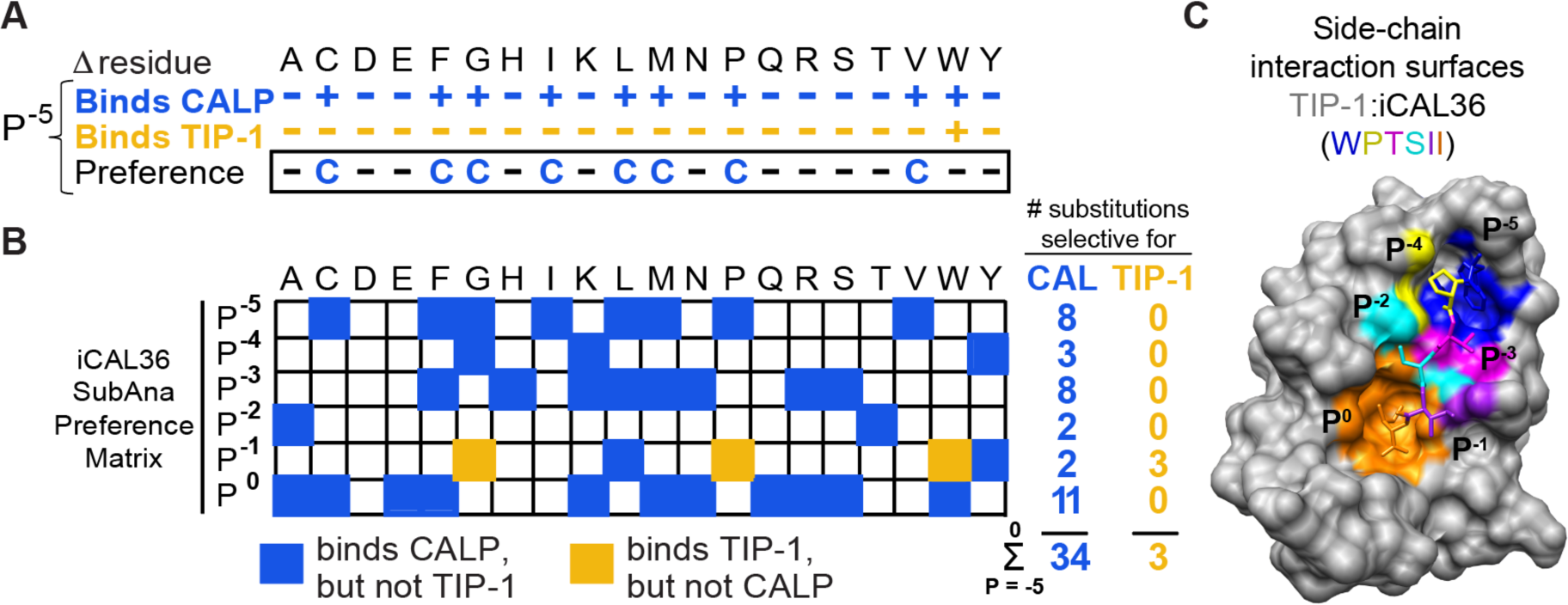
Using quantification of peptide array data reported here and previously [44], we identified amino-acid substitutions that retained ≥ 33% of wild-type sequence binding based on spot intensities. (**A**) Looking at substitutions in the iCAL36 sequence for the P^-5^ position, there are 9 amino acids, including Trp, that bind CALP, but only one, the wild-type Trp, that binds TIP-1. (**B**) Extending this analysis further, we quantified SubAna data for the C-terminal six positions of iCAL36. A blue box indicates a substitution, which may include the wild-type amino acid, that retains binding to CALP, but not TIP-1. A yellow box indicates a substitution that retains binding to TIP-1, but not CALP. The sum indicates there are 34 potential selective substitutions in iCAL36 that will bind CALP, but only 3 that will bind TIP-1. (**C**) The side-chain interaction surfaces for each iCAL36 position is shown on the TIP-1:iCAL36 structure. TIP-1 is in gray surface representation. The peptide is shown as sticks, with atoms colored by residue number, and the surface on TIP-1 that interacts with a given position colored accordingly. Here, P^0^ Ile = orange, P^-1^ Ile = purple, P^-2^ Ser = cyan, P^-3^ Thr = pink, P^-4^ Pro = yellow, and P^-5^ Trp = blue.

Overall, PDZ domain specificity can be described using neither degenerate nor highly differentiated motifs, but is instead a function of the overlapping core motif and modulatory selectivity determinants along the entire PDZ binding cleft (**Figure 6**). Furthermore, despite the overlapping nature of PDZ domain interactions, we argue that the relevant therapeutic window for targeting an individual domain can be defined in terms of selectivity differences and binding free energies, leading to the ability to engineer peptides with single-PDZ specificity in relevant cellular systems. The addition of *in vitro* techniques, e.g., the holdup assay, can additionally be utilized if necessary for pharmacological considerations, dependent on drug delivery method. Taken together, our work confirms that modulator preferences are important for PDZ network wiring, and additionally, can be exploited for therapeutic benefit.

## Material and methods

### Protein expression and purification

CALP (UniProt accession #Q9HD26-2) was expressed and purified as described previously [20,48,49,81]. TIP-1 (accession # O14907) was expressed and purified similarly except that an N-terminal His_10_ tag was used with a modified HRV 3C protease recognition sequence (LEVLFQ*G) upstream of the full-length protein sequence. Briefly, the TIP-1 PDZ domain was inserted into the pET16b vector (Blue Heron Technology, Inc.), and expressed in *Escherichia coli* BL21 (DE3) RIL cells. Following purification via immobilized metal-affinity chromatography, the protein was applied to a Superdex S75 gel filtration column (GE Healthcare) and eluted in gel filtration buffer [50 mM Tris pH 8.5, 150 mM NaCl, 0.1 mM TCEP, 0.02% NaN_3_]. Human rhinovirus 3C protease (Novagen) was added to the pooled protein fractions at a 1:30 mass ratio and incubated at 4 °C for 48 h. Following cleavage the protein was passed through a 1 mL HisTrap HP column (Cytiva) equilibrated in 20 mM imidazole, 25 mM Tris pH 8.5, 150 mM NaCl, 0.1 mM TCEP, 0.02% NaN_3_. The protein was further purified on a Superdex S75 column. Following gel filtration, the protein was dialyzed into gel filtration buffer, defined above, with 5% glycerol. TIP-1 protein was quantitated by using the A_280_ and the experimental extinction coefficient value of 10715 cm^-1^*M^-1^, determined as described [35].

### Peptide synthesis

All peptides except those used in peptide array experiments were synthesized and HPLC purified by the Tufts Peptide Core Facility. Fluorescein coupled to peptides via an N-terminal aminohexanoic acid linker is denoted by the prefix “*F\****-**”. Biotin coupled to peptides N-terminally is denoted by the prefix “*BT-*” where indicated. All sequence information is below.

### Cell culture

CFBE41o-cells [82] stably expressing *F508del-CFTR* under the control of a cytomegalovirus promoter (CFBE-ΔF cells) [83] were a generous gift of Dr. J.P. Clancy (University of Alabama, Birmingham). Cells were cultured as described [44] and switched to MEM containing only penicillin and streptomycin 24 h before experiments. All cells used in experiments were between passages 15 and 20.

### Fluorescence anisotropy and peptide array experiments

Peptide fluorescence anisotropy binding studies were performed and analyzed as described [20,35,44,45,49]. In the case of affinity, inhibitors with *K*_i_ > 1000 µM, affinity was described as undetectable. The following reporter peptides were used for competition *K*_i_ experiments: *F*\*-iCAL36 (*F*\*-ANSRWPTSII) for CALP (*K*_D_ = 1.3 μM) and TIP-1 (*K*_D_ = 0.54 μM); *F*\*-CFTR6 (*F*\*-VQDTRL) for N1P1 (*K*_D_ = 0.37 μM) and N1P2 (*K*_D_ = 1.08 μM); and *F*\*-CFTR10 (*F*\*-TEEEVQDTRL) for N2P1 (*K*_D_ = 0.32 μM) and N2P2 (*K*_D_ = 0.23 μM) [20,35,45,49]. Competition experiments were performed at protein concentration = 1.5\**K*_D_. Inverted peptide-array experiments were performed as described [45,80,84]. For TIP-1 peptide-array experiments, His_10_ tagged (uncleaved) protein was used to facilitate quantification.

### PDZ pull-down assays and mass spectrometry

Biotin-conjugated peptides or buffer were incubated with streptavidin paramagnetic beads (Promega). Excess peptide was removed by washing with wash buffer [50 mM Tris pH 8.0, 150 mM NaCl, 2 mM ethylenediaminetetraacetic acid (EDTA) pH 8.0, 2 mM ethylene glycol tetraacetic acid (EGTA) pH 8.0, 5% (*w/v*) glycerol, 1 mM dithiothreitol (DTT)]. Clarified CFBE-ΔF cell lysates (lysis buffer is wash buffer, plus one Complete tablet per 50 ml (Roche) and 1% (*w/v*) Triton X-100) were added to the beads and incubated with rotation for 90 minutes at 4 °C. Beads were washed and bound proteins were eluted with non-biotinylated cognate or scrambled peptide, sequences below. Proteins were separated by SDS-PAGE and immunoblotted as indicated. For preliminary mass spectrometry identification, SilverQuest (Invitrogen) was used to silver stain protein bands according to manufacturer’s instructions. Bands were considered to be candidate protein interactors if they were enriched in the specific non-biotinylated peptide-eluted lane (e.g. iCAL36) versus the SCR-eluted lane. Destained protein bands were submitted to Taplin (Harvard) or University of Vermont Proteomics facilities for identification.

The optimized pull-down and mass spectrometry protocol was slightly different, although the wash and lysis buffers were identical (see above). For each experiment, 1 mM peptide was used, instead of 0.5 mM. The sequences of the biotinylated peptides were: *BT*-iCAL36/*BT*-36 (biotin-WrFKKANSRWPTSII, where WrFKK is a linker sequence (r = D-Arg)), *BT*-L-iCAL36/*BT*-L-36 (biotin-WrFKKANSRLPTSII), *BT*-CFTR (biotin-TEEEVQDTRL), *BT*-HPV (biotin-WrFKKSSRTRRETQL), and *BT*-SCR (biotin-WrFKKSPTINSAIWR). The elution peptide sequences were: iCAL36 (ANSRWPTSII), L-iCAL36/L-36 (ANSRLPTSII), CFTR (TEEEVQDTRL), HPV16 (SSRTRRETQL), and SCR (SPTINSAIWR). Proteins were run to half the length of an SDS-PAGE gel, then stained using Bio-Safe Coomassie stain (Bio-Rad). Regardless of the appearance of distinct protein bands, the entire sample was cut into 4 sections, proteolyzed using trypsin (Promega), and analyzed by LC-MS/MS.

### LC-MS/MS Analysis

LC-MS/MS analysis was performed on an LTQ-Orbitrap mass spectrometer (Thermo Fisher Scientific, Bremen, Germany) equipped with an Agilent 1100 capillary HPLC, FAMOS autosampler (LC Packings, San Francisco, CA) and nanospray source (Thermo Fisher Scientific). Peptides were redissolved in 5% ACN / 1% formic acid and loaded onto an in-house packed polymer-fritted [85] trap column at 2.5 µl/min (1.5 cm length, 100 µm inner diameter, ReproSil, C_18_ AQ 5 µm 200 Å pore (Dr. Maisch, Ammerbuch, Germany) vented to waste via a micro-tee. The peptides were eluted by split-flow at ∼1000 psi head pressure from the trap and across a fritless analytical resolving column (20 cm length, 125 µm inner diameter, ReproSil, C_18_ AQ 3 µm 200 Å pore) pulled in-house (Sutter P-2000, Sutter Instruments, San Francisco, CA) with a 50 min gradient of 5-30% LC-MS buffer B (LC-MS buffer A: 0.0625% formic acid, 3% ACN; LC-MS buffer B: 0.0625% formic acid, 95% ACN). An LTQ-Orbitrap method consisting of one Orbitrap survey scan (AGC Orbitrap target value: 700K; R = 60K; maximum ion time: 800 milliseconds; mass range: 400 to 1400 *m/*z; Orbitrap “preview” mode enabled; lock mass [86] set to background ion 445.120029) was collected, followed by ten data-dependent tandem mass spectra on the top ten most abundant precursor ions (isolation width: 1.6 *m/*z; CID relative collision energy: 35%; MS1 signal threshold: 12,500; AGC LTQ target value: 3,500; maximum MS/MS ion time: 125 milliseconds; dynamic exclusion: repeat count of 1, exclusion list size of 500 (max), 24 seconds long, +/- 20 ppm, and doubly- and triply-charged precursors only).

### Peptide spectral matching and bioinformatics

Raw data were searched using SEQUEST [87,88] (ThermoFisher Scientific, San Jose, CA) against a target-decoy (reversed) [89] version of the human proteome sequence database (UniProt; downloaded 9/2013; 74,338 total (forward & reverse) proteins) with a precursor mass tolerance of +/- 1 Da and requiring fully tryptic peptides with up to two mis-cleavages, carbamidomethylcysteine as a fixed modification and oxidized methionine. The resulting peptide spectral matches were filtered to < 1% false discovery rate (FDR), based on reverse-hit counting.

Proteins were considered identified if ≥ 2 constituent peptides were sequenced. PDZ domain-containing proteins that were present in all samples, including the negative control (*BT-*SCR) included PDLI1, ZO2, LMO7, ZO1, and PDLI7. PDZ domain-containing proteins for which only 1 peptide was identified included: AHNK (*BT*-SCR, *BT*-CFTR), DLG1 (*BT*-iCAL36, *BT*-CFTR), GIPC1 (*BT*-L-iCAL36), LIN7A (*BT*-HPV), LNX2 (*BT*-iCAL36), MPDZ (*BT*-iCAL36), NHRF2 (*BT*-iCAL36), PDLI3 (*BT*-L-iCAL36, *BT*-CFTR), SNTB2 (*BT*-iCAL36, *BT*-L-iCAL36), SNX27 (*BT*-CFTR), TIP-1 (*BT*-CFTR).

### Crystallization and structure determination

Peptides (iCAL36, F-iCAL36, Y-iCAL36) were added at a final concentration of 1-5 mM to purified TIP-1 at 5.5 mg ml^-1^ in 10 mM HEPES pH 7.4, 25 mM NaCl. Crystallization conditions identified by screening were optimized in hanging-drop format at 291 K, by adding 2 µl of the complex in screening buffer to 2 µl reservoir solution. The reservoir contained 500 µl solution. Crystals appeared in 2-4 days and continued to grow for up to 14 days. The crystals used for data collection were obtained using: for iCAL36 (1 mM peptide used), 100 mM NH_4_SCN, 100 mM MES pH 6.0, 36% (w/v) polyethylene glycol (PEG) 1000 as reservoir buffer; for F-iCAL36 (up to 5 mM peptide), 200 mM NH_4_SCN, 100 mM sodium acetate pH 5.0, 31% (w/v) PEG 4000; and forY-iCAL36 (up to 5 mM peptide), 150 mM KSCN, 100 mM MES pH 6.0, 40% (w/v) PEG 3350.

For data collection, a crystal was transferred into cryoprotectant buffer [150-200 mM NH_4_SCN (or KSCN), 100 mM MES pH 6.0, 30-50% (w/v) PEG 400]. The data sets used for structure determination were obtained at 100K, λ=1.0000 Å on beam line X6A at the National Synchrotron Light Source (NSLS) at Brookhaven National Laboratory. For TIP-1:iCAL36, two data sets were collected and later merged; one over a 360° range, using 0.3° frames and an exposure time of 2 s per frame, and the second over 60°, using 1° frames, an exposure time of 0.5 s per frame, and an aluminum attenuator. For TIP-1:F-iCAL36 and TIP-1:Y-iCAL36, single data sets were collected at Δφ=0.3° over 360°. Diffraction data were collected to high (1.24, 1.5, and 1.6 Å, respectively) resolution and processed using the XDS package [90,91]. Data collection and refinement statistics are presented in **Table 2**.

Molecular replacement was performed using PHENIX [92], using the TIP-1 PDZ domain from PDB entry 3DIW as a template [74]. The model was built and refined using PHENIX. In all cases, electron density for the peptide prior to its inclusion in the model confirmed its presence and served as a template for initial model building (**Figure S2**). Following refinement, model geometry was assessed using MOLPROBITY and the PDB validation server [93].

### Ussing chamber measurements

Short circuit current (I_SC_) measurements were performed as described [44]. Briefly, for Ussing chamber measurements, 10^5^ cells were seeded onto 12 mm Snapwell permeable supports (Corning) and allowed to form polarized monolayers over the course of 9 days. CFBE-ΔF cells were dosed with 0.5 mM peptide via BioPorter (Sigma) 3.5 h before the start of Ussing chamber measurements. Cells were maintained at 37 °C throughout treatments; the DMSO concentration did not exceed 0.03%. Cells were treated sequentially with 50 µM amiloride, 20 µM forskolin, 50 µM genistein, and 5 µM CFTR_inh_-172 in 5.0 min intervals. CFTR-specific chloride efflux was computed as the magnitude of ΔI_SC_ following application of CFTR_inh_-172. Resistances were monitored throughout each experiment to ensure monolayer integrity.

### Statistical Analysis

Values are reported as mean ± SD except for Ussing-chamber experiments where mean ± SEM is reported. Student’s one-tailed t-test was used for fluorescence anisotropy binding experiments while the Student’s one-tailed paired t-test was used for analysis of Ussing-chamber experiments.

## CRediT authorship contribution statement

**Jeanine F. Amacher**: Conceptualization, Data curation, Formal analysis, Investigation, Validation, Visualization, Writing – original draft, Writing – review & editing. **Patrick R. Cushing**: Conceptualization, Data curation, Formal analysis, Investigation, Validation, Visualization, Writing – original draft, Writing – review & editing. Lars Vouilleme: Formal analysis, Investigation, Writing – review & editing. **Sierra N. Cullati**: Formal analysis, Investigation, Writing – review & editing. **Bin Deng**: Investigation, Methodology, Writing – review & editing. **Scott A. Gerber**: Methodology, Resources, Supervision, Writing – review & editing. **Prisca Boisguerin**: Funding acquisition, Methodology, Resources, Supervision, Writing – review & editing. **Dean R. Madden**: Conceptualization, Funding acquisition, Project administration, Resources, Supervision, Data curation, Formal analysis, Methodology, Validation, Visualization, Writing – original draft, Writing – review & editing.

## Acknowledgments

We thank Dr. Justin Piro for expression and purification of NHERF2 PDZ2, Dr. Devin Schweppe for helpful scientific discussion, Bonita Coutermarsh and Roxanna Barnaby for cell-culture technical support and Dr. Vivian Stojanoff, Dr. Jean Jakoncic, and Edwin Lazo (NSLS) for assistance with X-ray data collection and interpretation. We also gratefully acknowledge the collaborative support of Dr. B. Stanton (Dartmouth), Dr. W. Guggino and Dr. J. Cheng (Johns Hopkins University), and Dr. R. Volkmer (Charité Universitätsmedizin). This research project was supported by NIH grants R01-DK101451, P20-GM113132, P30-DK117469, P30-GM106394, and T32-GM008704 (JFA, PRC, SNC, DRM), the Cystic Fibrosis Foundation Research Development Program (STANTO19R0), and by Mukovizidose e.V. (S05/08), the German Cystic Fibrosis Association (LV, PB). The VBRN Proteomics Facility is supported through NIH grant P20-GM103449. Beamline X6A at NSLS I was funded by the NIH NIGMS (GM-0080) and the US Department of Energy (No. DE-AC02-98CH10886).

