## Supplementary material for "Sequence engineering at non-motif modulator residues yields a peptide that effectively targets a single PDZ protein in a disease-relevant cellular context"

by

Jeanine F. Amacher, Patrick R. Cushing, Lars Vouilleme, Sierra N. Cullati, Bin Deng, Scott A. Gerber, Prisca Boisguerin, and Dean R. Madden

### Table of Contents

**Table S1. Identified proteins in pull-down with mass spectrometry experiments.** 2

**Figure S1. iCAL36 interacts with TIP-1 in airway epithelial cell lysates.** 3

**Figure S2. Quality of omit and final electron density maps of iCAL36 Trp side chain.** 4

**Figure S3. TIP-1 structures reveal canonical Class I PDZ binding and accessibility of biotin in CAL PDZ:*BT*-L-iCAL36 structure.** 5

**Figure S4. CombLib peptide arrays ($P^{-5}$, $P^{-4}$) for CAL and TIP-1.** 6

**Figure S5. A 'therapeutic window' for selective CALP peptide inhibition.** 7

**Table S1. Identified proteins in pull-down with mass spectrometry experiments.**

| UniProt ID | Abbreviation | Protein Name |
|---|---|---|
| GOPC_HUMAN | CAL | Golgi-associated PDZ and coiled-coil motif-containing protein |
| CSKP_HUMAN | CASK | Calcium/calmodulin-dependent serine protein kinase |
| DLG1_HUMAN | DLG1 | Disks large homolog 1 |
| INADL_HUMAN | INADL | InaD-like protein |
| LIN7C_HUMAN | LIN7C | Protein lin-7 homolog C |
| MPP2_HUMAN | MPP2 | Membrane-associated guanylate kinase (MAGUK) p55 subfamily member 2 |
| PALS1_HUMAN | MPP5 | Membrane-associated guanylate kinase (MAGUK) p55 subfamily member 5 |
| PALS2_HUMAN | MPP6 | Membrane-associated guanylate kinase (MAGUK) p55 subfamily member 6 |
| NHRF1_HUMAN | NHERF1 | Na(+)/H(+) exchange regulatory factor 1 |
| NHRF2_HUMAN | NHERF2 | Na(+)/H(+) exchange regulatory factor 2 |
| NHRF3_HUMAN | NHERF3 | Na(+)/H(+) exchange regulatory factor 3 |
| SCRIB_HUMAN | SCRIB | Scribble 1 homolog |
| SNTB2_HUMAN | SNTB2 | Beta-2-syntrophin |
| TX1B3_HUMAN | TIP-1 | Tax1-binding protein 3 |

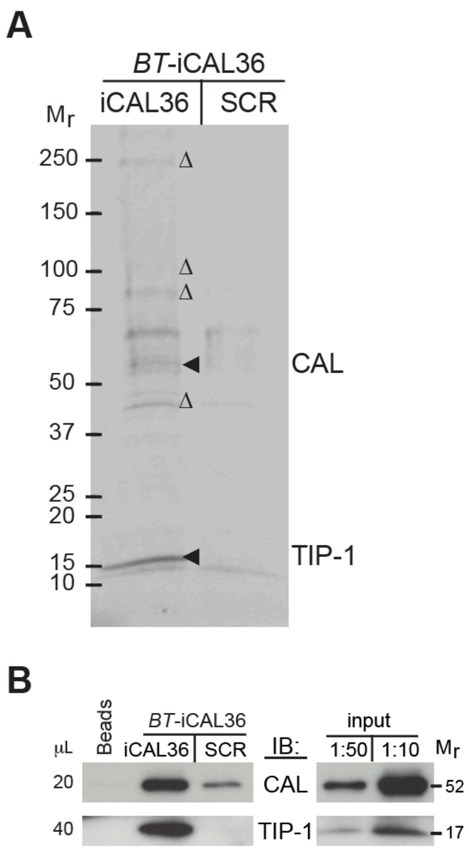


**Figure S1. iCAL36 interacts with TIP-1 in airway epithelial cell lysates. (A)** Endogenous proteins were captured from CFBE-ΔF lysates using immobilized *BT*-iCAL36 as bait, eluted with iCAL36 or a scrambled control peptide (SCR) and visualized by SDS-PAGE. A representative gel is shown (n=3). **(B)** α-CAL or α-TIP-1 immunoblots (IB) confirm the enrichment of endogenous CAL and TIP-1 proteins in iCAL36 pulldowns.

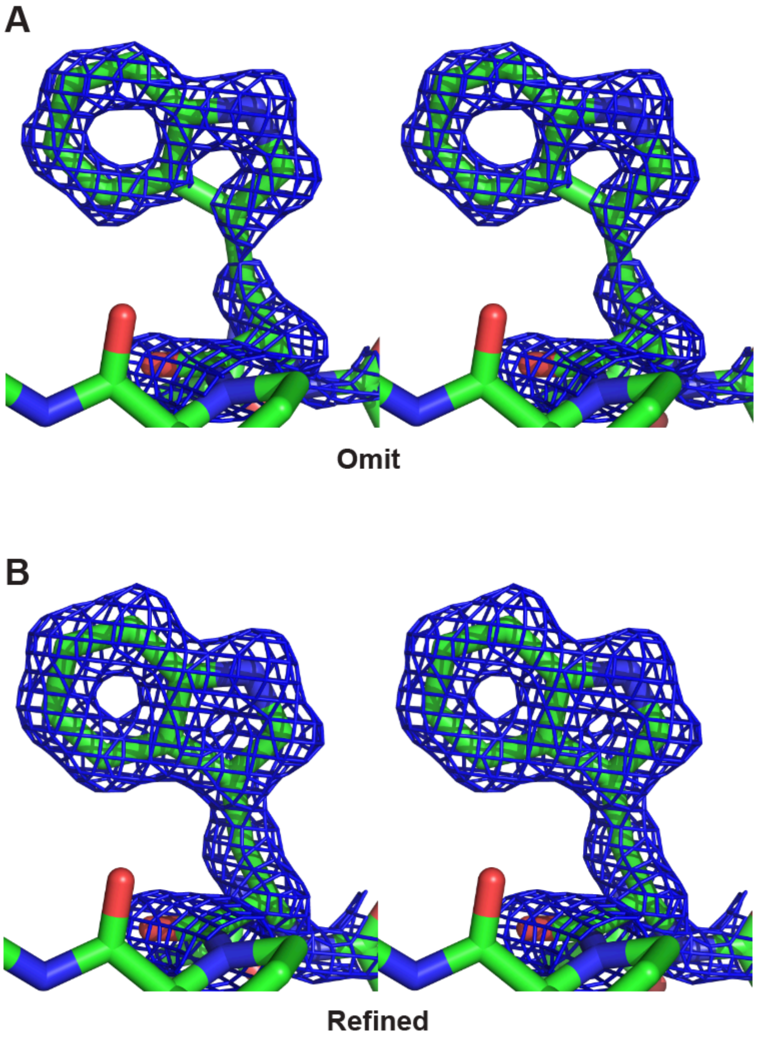


**Figure S2. Quality of omit and final electron density maps of iCAL36 Trp side chain.** (**A**) The P-5 Trp side chain of the refined iCAL36 peptide model is shown together with electron density calculated prior to inclusion of peptide residues in the model. The omit map is contoured at 1.5σ, and shows unambiguous tryptophan density at the correct location. (**B**) The peptide model is shown together with the electron density computed in the final round of refinement. The refined map is contoured at 1.5σ.

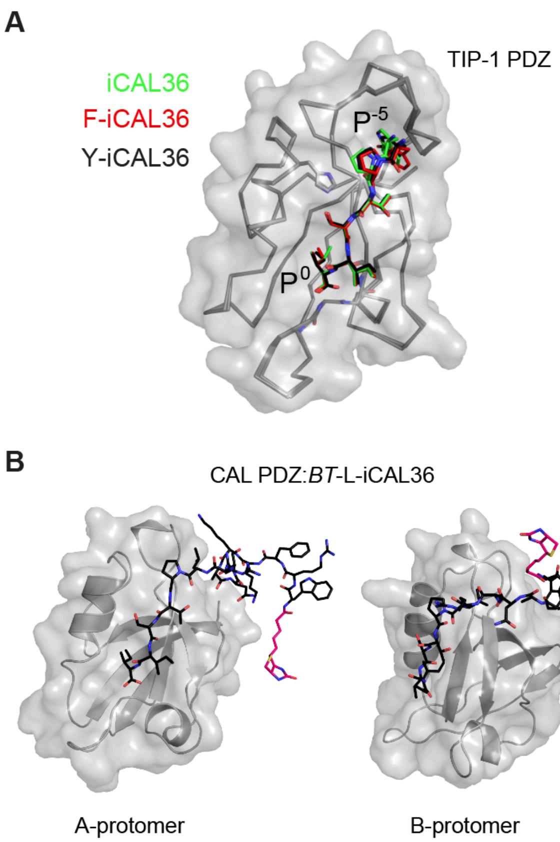


**Figure S3**. **TIP-1 structures reveal canonical Class I PDZ binding and accessibility of biotin in CAL PDZ:*BT*-L-iCAL36 structure. (A)** The structures of the TIP-1 (grey, surface representation and $C_\alpha$ trace) with the iCAL36 (green), F-iCAL36 (red), and Y-iCAL36 (black) peptides (stick figure) reveal a canonical PDZ binding interaction involving residues $P^0$ through $P^{-5}$. **(B)** The biotin moiety (hot pink carbon atoms, sticks) is accessible in the A-protomer (left) of the CAL PDZ:*BT*-L-iCAL36 structure (grey, surface and cartoon representation). In the B-protomer (right), the biotin is less accessible, but this could be due to differential crystal packing between the two protomers. In both cases, the most substantial interactions with the peptide (black carbon atoms, sticks) are at residues P0 to $P^{-5}$ position, consistent with previously determined structures. Atoms in the peptide are colored by element (red=O, blue=N, yellow=S).

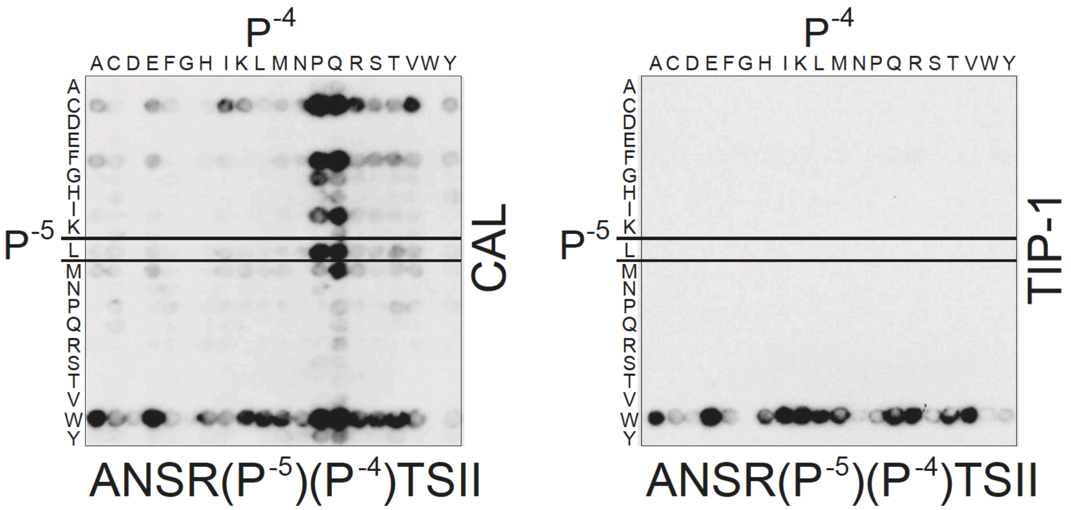


**Figure S4. CombLib peptide arrays ($P^{-5}$, $P^{-4}$) for CAL and TIP-1.** CombLib peptide arrays varying two positions ($P^{-5}$, $P^{-4}$) of the iCAL36 sequence were incubated with either CAL (left) or TIP-1 (right) PDZ domains. Only peptides with a Trp residue at $P^{-5}$ bound TIP-1, including iCAL36 (WP). Incubation with CAL shows higher redundancy at this position. Of peptides with Pro at $P^{-4}$, those with Leu, Phe or Cys at $P^{-5}$ bound CAL most strongly, but did not bind TIP-1.

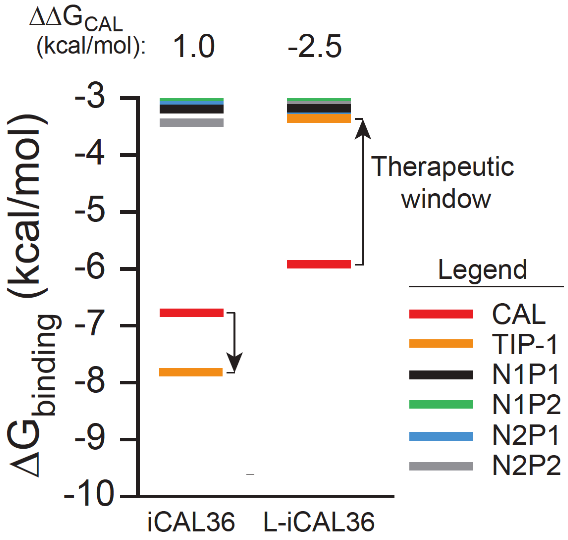


**Figure S5. A 'therapeutic window' for selective CALP peptide inhibition.** $\Delta G^0$ values calculated from fluorescence anisotropy competition experiments are shown for the indicated peptide sequences against NHERF1, NHERF2, CAL, and TIP-1 PDZ domains. The $\Delta\Delta G$ value from CALP to the highest affinity off-target interaction is indicated by an arrow for each sequence.
